# Viscous deformation of relaxing ventricle and pulsatile blood propelling (numerical model)

**DOI:** 10.1101/009522

**Authors:** Yuri Kamnev

## Abstract

The numerical model of one-loop circulation exploits viscous deformation as mechanism of ventricular filling. Mathematical advantage of viscous deforming is a possibility to present the ventricular filling as the function of two independent variables (stress and time); two-dimensional argument frames a table which permits to calculate end-diastolic ventricular volume due to information about measured venous pressure and duration of ventricular diastole. The equation was deduced which balances the system while varying of such parameters as arterial resistance, the value of normal rhythm and the volume flow rate. The model pays attention on the phenomenon of asymmetrical position of normal rhythm (the steady rhythm in conditions of rest) and explains why the operating range of brady-rhythms is much narrower than the operating range of tachy-rhythms.

## Introduction

The opinion that ventricular relaxation is a viscoelastic process can be found mentioned [1] but complexity of material adds nothing comprehensible for connection of such a specific behavior of material to the controlling of circulatory parameters. Kelvin-Foight model is putting in action the viscous and the elastic processes in parallel; it results in the delayed reaction of the elastic material (so called, “creep”) which gives no development of understanding the ventricular filling. Maxwell model of complex material, which combines the two components operating in series, can be considered justified if the elastic component plays, for example, the role of safety device preventing the excessive dilatation. Nevertheless the operating range of dilatation (i.e., the range of filling itself) needs a description in terms of isolated viscous deforming because it possesses two advantages which ought not to be contaminated by combination with elastic deformation. Firstly, as far as viscous deformation demonstrates the behavior of liquid all the information about the ventricular filling with blood (liquid) transmits to the relaxing myocardium if it also behaves like liquid (imitates the behavior of liquid during relaxation). Secondly, viscous deformation depends on the duration of process (unlike elastic one) and, consequently, pulsatile mode of propelling of blood, - which proposes invariable duration of systole and changeable duration of diastole, - must have affinity to the time-dependent process of filling because in that case the linkage between deformation and rhythm appears (and, consequently, deformation becomes controllable). Simple numerical model of one-loop circulation that is presented here solves the problems of balancing of several basic physiological parameters in response to deviations of the system. Finally the model explains why the normal rhythm (steady rhythm in the conditions of rest) is located asymmetrically between operating range of brady-rhythms (which is quite narrow) and operating range of tachy-rhythms (which is much broader).

## Methods

The general definition of viscous deformation is the following: tensor of stresses is a linear function of tensor of strain rates of elementary volume of liquid. We may use more simple definition (Newton's law for internal friction): 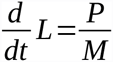, where *L* is a deformation, *P* is a pressure (stress) and *M* is the coefficient of dynamic viscosity with dimensions of quantity 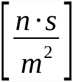; hence, 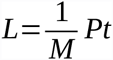 is the formula simplified for arithmetic calculations. *L* is the relative viscous deformation and has no dimensions of quantity (abbreviated by cancellation) 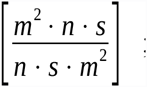, if we want to operate with absolute viscous deformation (*L* gains the dimensions of volume 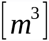) the coefficient of dynamic viscosity *M* must be changed in respect of its dimensions of quantity from 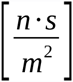 to 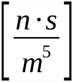 and its physical interpretation alters either. Absolute viscous deformation *L* 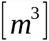 depends on 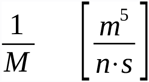 and, consequently, *L* will grow not only due to the lower viscosity of liquid but also due to the larger volume of empty room which the definite geometry of space can place at our disposal; in other words, we must take into consideration that some outer restriction can produce the correction of the volume that we are expecting to observe according to the law of viscous deformation which performs at some ideal geometry of space.

The comprehension of this relationship is more clear in terms of hydrodynamics. The parameter with dimensions of quantity 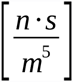 corresponds to hydrodynamic resistance *R* determined by standard equation 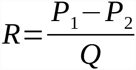, where *Q* is a volume flow rate and *P*_1_–*P*_2_ is the difference of pressures at the beginning and at the end of the pipe. We may represent *Q* as 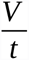, where *V* is a volume and *t* is a time, and then rewrite hydrodynamic equation: 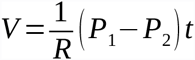. Compare it with the law of viscous deformation 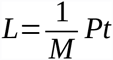 and the interpretation of 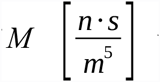 will be readable due our understanding of hydrodynamic resistance 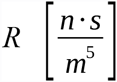. The resistance is the compound parameter that includes either viscosity of the liquid or the friction over the wall of the pipe and the length and diameter of the pipe. All characteristics of the pipe can be called the restrictions originated from definite spatial geometry. Therefore, when we use coefficient 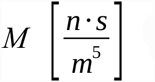 we operate not only with viscosity but also with some spatial restrictions - external conditions, or medium, where the viscous deformation takes place.

For simplification of modeling the dynamical viscosity itself can be approximated to the viscosity of water (*M* = 1 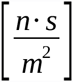 at temperature 20° C), and, respectively, the “external restrictions” - i.e., the another component of 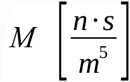, - can be assumed negligible. Therefore, we have clarified the position concerning the parameters we are dealing with: firstly, it is the absolute viscous deformation *L* 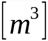 and, secondly, it is the coefficient 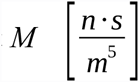 with physical interpretation different from the coefficient of dynamic viscosity 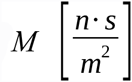, although numerically we assume both coefficients equal.

Thus, *L* is the ventricular volume which is enlarging during ventricular diastole, *P* denotes the venous pressure, *t* – the duration of ventricular diastole, and 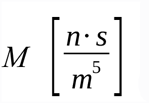 is the coefficient we have discussed above. Viscous deformation permits to construct a table – a two-dimensional structure, - because viscous deformation depends on two variable parameters, *P* and *t* (i.e., stress and duration of stress); viscous deformation differs fundamentally from elastic deformation which depends on the value of stress only and is independent of the duration of stress. The table at **Fig.1** is framed on the basis of formula 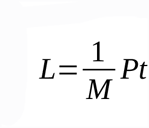 described above.

**Fig.1.**
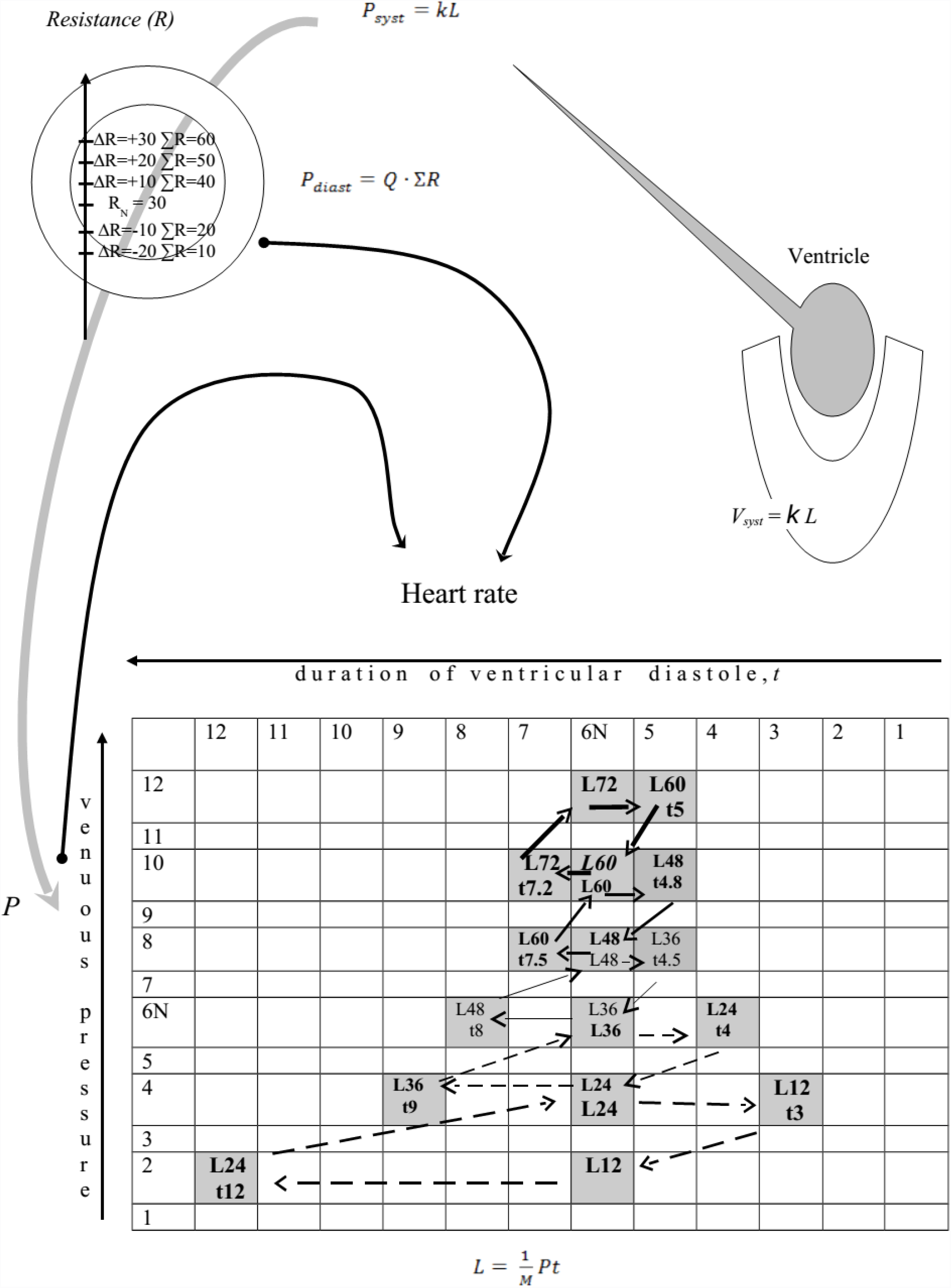
End-diastolic deformation of the ventricle (*L*) is generated by two variables due to the law of viscous deforming – venous pressure (*P*) and duration of ventricular diastole (*t*). Frank-Starling law converts end-diastolic volume into systolic volume which is assumed numerically equal to systolic pressure that originates hydrodynamics in the pipe according to formula *P*_*sys*_ – *P* = *Q · ΣR*, where *Q* is a volume flow rate in the arterial pipe, ∑*R* is a resistance proceeded from the work of arterial resistive vessels (*Q*·*ΣR* can be interpreted as diastolic pressure). Two reflexes (black curves begin from zones of measuring of pressure, - arterial or venous, - and both end at pace maker) regulate the responses. The scale of variation of ∑*R* is shown; different values of ∑*R* effect the correction of *t* jointly with *P* according to the deduced equation (see the text); line№6 and column№6 denote normal venous pressure and normal duration of ventricular diastole, respectively. Each type of arrows forms a circle which depicts deviation and restoration of the system at each level of ∑*R*.

The calculated deformation *L* is the end-diastolic volume which is directly proportional to the systolic volume due to Frank-Starling law [2], i.e., *V*_*syst*_ = *k L*, where *k* is a coefficient of proportionality without dimensions of quantity.

Proportionality of volumes found experimentally is quite close to the function of direct proportionality *V*_*syst*_ = *f* (*V*_*diast*_) with coefficient equal to 1. It differs from the linkage between contractile force, *F*, and linear elongation of muscular fiber, *l*, according to the function *F* = *f*(*l*^2^) that was deduced theoretically [3] and that was demonstrated by experimental curves looking very much like the ascending limb of parabola. Such portion of ascending limb of parabola can be linearized by approximation to direct proportionality (with the slope of line corresponding to coefficient more then 1 and with a shift along abscissa axis). The domain that organizes the above portion of parabola reflects the restricted elongation of fiber. Variation of diastolic volume at *V*_*syst*_ = *f*(*V*_*diast*_) possesses the analogous restriction of variation; consequently, the linearized values of contractile force from *F* = *f*(*l*^2^) and the range of systolic volume from *V*_*syst*_ = *f*(*V*_*diast*_) may be considered linked by some linear operator. We may assume the steady work of such operator and use phenomenological relationship *V*_*sys*_ = *f* (*V*_*diast*_) at our speculations instead of fundamental law *F* = *f*(*l*^2^).

The significant pressure at the initial portion of arterial pipe, - the portion which is located between the ventricle and arterial resistive vessels and is presented by elastic artery with constant resistance (neglected at the present model), - exists only during the period of ejection of blood from the ventricle. In other words, we assume that the pressure, at this portion of pipe, has a significant value only during ventricular systole but has a negligible value during ventricular diastole. Factual appearance of non-ignored pressure at this portion of pipe, i.e., diastolic pressure, is a consequence of a liquid to conduct pressure, - but this pressure (diastolic pressure) has been generated not at the initial portion of pipe. Therefore, at the initial portion of pipe we observe during the diastole the superimposition of a negligible value of pressure (which is also essential to this portion of pipe after the cessation of ejection) and some substantial value of pressure which has been conducted from the another portion of pipe where this substantial pressure has the origin of generation. It is a first application of the approach we suggest as a method of simulation of pulsatile blood flow; the principle is the following: it is necessary to combine three parameters (to construct the triple parameter concerning pressure):

- the anatomical region of a pipe (i.e., we substitute the sequence of regions instead of flow travel, i.e., we simulate a movement);
- the relation to the beginning, systole, or to the end, diastole, of cardiac cycle (i.e., we simulate the pulsation);
- the pressure.

Before we quit the initial portion of pipe (with significant pressure attached to the systole) let us determine the linkage of this pressure (systolic pressure) with the mechanism of its generation. Frank-Starling law places the systolic volume at our disposal and it is necessary to convert it into systolic pressure. Within present modeling we assume the transition from systolic volume to systolic pressure a numerical equalization but the dimensions of quantity must be specified. Consequently, if *V*_*syst*_ = *k L and k* is a coefficient of proportionality without dimensions of quantity we need to change the coefficient in *P*_*syst*_*=kL*, where *k* now is a coefficient of proportionality with dimensions of quantity 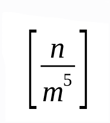. Utmost simplification claims that *k* is equal to 1, and, therefore, we assume that systolic pressure is numerically equal to end-diastolic deformation. (As for conversion of systolic volume into systolic pressure the above numerical equalization is not absolutely unfounded. At least the linearity of linkage of systolic volume and systolic pressure can be demonstrated by application of the present model of circulation to some observed physiological phenomenon [4]).

Let us continue attaching pressures to the anatomical portions of pipe and to cardiac cycle. The formula that determines the bond between the pressure difference at the beginning and the end of a pipe, *P*_1_*–P*_2_, the volume flow rate, *Q*, and the hydrodynamic resistance, *R*, is standard 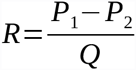. It can be represented as *P*_*syst*_–P = *Q*· *ΣR*, where *Q*· *ΣR* has the dimensions of quantity of pressure and so let us associate this pressure with the pressure that we observe during the late period of cardiac cycle (diastole) and let us assume that the diastolic pressure can be registered, as the essential parameter, only at the basin of arterial resistive vessels; therefore, we speak about diastolic pressure: *P*_*diast*_ = *Q*· *ΣR*. Diastolic pressure will inevitably be conducted from the basin of resistive arteries to the initial portion of the pipe where the essential pressure during the diastole is negligible.

The final portion of the pipe, - the vein, - possesses the pressure that can be called the residual pressure *P=P*_*syst*_–*P*_*diast*_ = *P*_*syst*_*-Q*· *ΣR* and this pressure is attached neither to systole nor to diastole (or is attached to both of them) due to the introduction of capillary damper which is located after the basin of resistive arteries. The resistance of capillary damper can be considered constant and at the present model we assume it ignorable; we remind that the resistance of the initial artery is neglected also and, consequently, the general resistance of the whole pipe consists only of the arterial resistance originated by resistive arteries, and this resistance is changeable and controllable.

We may interpret diastolic pressure as a subtrahend that takes away some quota of pressure from initial (systolic) pressure, i.e., it converts systolic pressure into venous pressure: *P*_*syst*_–*P*_*diast*_ = *P*, but simultaneously the subtrahend itself (diastolic pressure) has been generated from the initial pressure (systolic pressure) during the operation of subtraction. Anatomically the subtraction takes place at the region of resistive arteries but it is pertinent to emphasize that we can not speak about systolic pressure inside the region of resistive arteries as the essential pressure. The observed wave of elevated pressure which chronologically coincide with systole at the region of resistive arteries must be considered the pressure conducted from initial artery. Situation is symmetrical to the one at the region of initial artery where diastolic pressure is not essential but is the conducted pressure. Nevertheless, the region of the initial artery is the only portion of pipe where we can observe minuend (essential systolic pressure) and subtrahend (conducted diastolic pressure) separately, i.e., out of operation of subtraction. On the contrary, the region of resistive arteries produces the process of subtraction itself while generating of the subtrahend (diastolic pressure essential for this region) from the minuend (conducted systolic pressure which is losing its energy while the conversion into the subtrahend and the remainder).

It was mentioned that at the present model the capillary damper does not increase the resistance but only liquidates the division “systole-diastole”; therefore, the remainder (venous pressure) appears behind the operator of subtraction and is associated with the final portion of pipe (vein) supplemented with the lack of pulsatile flow.

Besides, the subtrahend consists of two components (∑*R* and *Q)* and it permits us to be aware of the volume flow rate if we know the value of diastolic pressure (pressure at the end of cardiac cycle which can be measured by baroreceptors in resistive arteries) and if we also possess the direct information from muscles of resistive arteries about the extent of narrowing of the pipe: 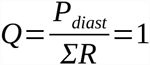. For the most part of further variations we consider *Q* of a constant value and equal to 1; in other words, the information from baroreceptors and the information from muscles of the resistive arteries will coincide. Such a condition is not groundless as far as it reflects the existence of some optimal capillary flow which corresponds to stability of biochemical processes of cell metabolism. Nevertheless, it does not mean that *Q* is not able to be increased, and such deviation will also be carried out and scrutinized at the final part of testing of the model. Therefore, ∑*R* is numerically equal to diastolic pressure when *Q* = 1, i.e., 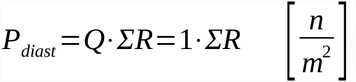.

The quanta of columns and lines of the table at **Fig.1** were arbitrary chosen but it was convenient to frame 12×12-table for making use of minimal whole numbers denoting increments of venous pressure and duration of ventricular diastole (although later, while deviating the system, the advantage of making operations with whole numbers will be lost); the normal value of venous pressure and the normal value of the duration of ventricular diastole were placed in the middle of the scales (line N<u>o</u>*6* and column N<u>o</u>6, respectively). Now it is necessary to determine the state of equilibrium based on normal parameters of: 1) venous pressure, 2) duration of ventricular diastole and 3) the resistance. Since *P=P*_*syst*_*–Q·ΣR* and 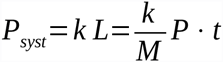, it is convenient to assume that normal resistance, *R*_N_, is equal to some number of 30 because it permits to get the remainder (normal venous pressure) equal to 6 after the subtraction of the value of diastolic pressure (*R*_N_ multiplied by *Q* = 1) from the value of normal ventricular end-diastolic deformation (which is numerically equal to systolic pressure). This quantity of 6 (normal venous pressure) participates in calculation of normal end-diastolic deformation – which is the product of normal venous pressure and normal duration of ventricular diastole, - and if we choose quantity of 6 either for the normal value of the duration of ventricular diastole (and taking into account that *M* is equal to 1 and *k* is equal to 1 either) we get 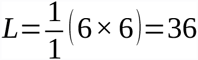. After subtraction of 30 (*P*_*diast*_*=Q · R*_N_=1 · 30 = 30) from 36 (*P*_*syst*_ is numerically equal to *L*) we get the quantity equal to 6. This quantity of 6 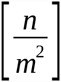 turns back to the table as the number of line that denotes the normal venous pressure *P*_N_ = 6. The numerical expression of equilibrium of three parameters is found and this equilibrium characterizes the normal, non-deviated circulation (one-loop circulation).

Two reflex arcs carry out the controlling and both begin from the reception of pressure: the one (analog of carotid reflex) decelerates rhythm in response to elevation of diastolic pressure, *P*_*diast*_, and accelerates rhythm in response to descending of *P*_*diast*_; this reflex measures pressure inside resistive artery at the end of cardiac cycle (diastolic period), - although we have mentioned that in general case the direct information from the muscles of resistive arteries is indispensable, i.e., the parallel pathway transporting such information must exist; the other reflex (analog of Bainbridge reflex which existence is hardly proved but the term is eligible) accelerates rhythm in response to elevation of venous pressure, *P*, and decelerates rhythm in response to descending of *P.*

## Results

Let us trace the reorganization of the system after some minimal, additional to normal, heightening of the resistance, ∆*R* = 10; hence, the new, i.e., deviated, value of the resistance is the following: *ΣR=R*_*N*_*+∆R* = 30 + 10 = 40. The elevated resistance needs the augmentation of contractile force, and the increasing of systolic pressure, respectively, - because the previous one, *P*_*syst*_ = 36, will not overcome the new value of diastolic pressure, *P*_*diast new*_:

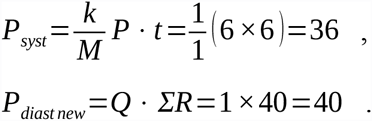

The only way to achieve the prevalence of *P*_*syst new*_ over *P*_*diast new*_ is to create higher value of systolic pressure my means of elongating of the duration of ventricular diastole, *t*, because the venous pressure is still normal, *P* = 6, and there is no direct influence upon it; moreover, we do not determine the duration of the delay of changing of venous pressure – the delay that inevitably appears due to the work of capillary damper.

So it is necessary to introduce one more assumption (condition) pertinent for this model: the reaction of carotid reflex upon the change of the resistance is extremely quick (within one heart beat); this assumption is indispensable for conservation of the value of venous pressure that was registered just before the resistance inside the arterial basin has changed. We substitute extreme quickness of the reaction instead of determination of the delay of change of venous pressure in response to the events happening inside the arterial basin – variations of the resistance (coinciding with diastolic pressure when *Q = 1*) and systolic pressure. In other words, the degree of inertness of converting of systolic pressure into venous pressure, -i.e., the speed of subtraction of *P*_*diast*_ from *P*_*syst*_ supplemented with liquidation of division “systole-diastole”, – i.e., factually the inertness generated mostly by capillary damper, - is omitted in the model.

Thus, quick reaction of carotid reflex is the following: the detection of the rise of *P*_*diast new*_ results in deceleration of rhythm. But what will be the new value of rhythm?

The answer needs some speculations. Variation of rhythm can be equalized (with inversion) to the variation of the duration of ventricular diastole because the duration of ventricular systole can be considered constant at any rhythm. Elongation of the ventricular diastole, – multiplied by the value of present venous pressure (which is considered normal), – will result in expanding of deformation; the expanded deformation will generate augmented contractile force (systolic pressure, respectively) which will permit to overcome the elevated resistance ∑*R* = 40. If the residual pressure (venous pressure) is still *P* = 6 (i.e., *P*_*syst new*_*—P*_*diast new*_ = *P*_*syst new*_*—Q* · *ΣR* = *P*_*syst new*_–1 × 40 = 6) it means that the mechanism of response is aimed to maintain venous pressure at constant value; in other words, the mechanism of response involves only the one variable parameter, – the duration of ventricular diastole, - into the process of expanding of viscous deformation, and the another parameter, venous pressure, remains not involved. Obviously such an approach leads to early depleting of the ability to react: few steps of the resistance rising will lead to the unreal values of deceleration of the heart rate. On the contrary, the approach with both parameters participating in expanding of deformation permits to reorganize circulation in response to broader diapason of rising and falling of the resistance. Therefore, the residual pressure (venous pressure) must be found on some higher level (7 or higher, for the present instance). Certainly the Bainbridge reflex will respond to; firstly, the reflex will detect the elevation of the venous pressure and, secondly, it will accelerate rhythm. The product of new *P*, which is higher then normal, and newly accelerated rhythm (newly shortened t), - i.e., the product that creates end-diastolic deformation of ventricle (which then will be transformed by the work of myocardium into systolic volume), - is aimed to retain the chosen higher level of venous pressure but we still have no criterion concerning what is “higher”. We may only assume that this higher level must be the nearest one unless we lose all the advantage of the second parameter – but what is the nearest level? Or maybe there is the optimal new level? At least we may frame the equation based on the following equalities:

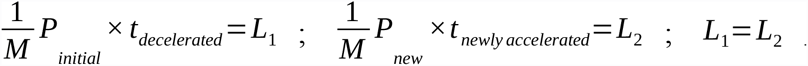

We considered *L*_1_ equal to *L*_2_ because we deal with the same deformation – the one is achieved by elongation of diastole (*P* is constant) and another is achieved by new higher level of *P* combined with restoration of initial duration of diastole which was registered before the elevation of the resistance and which we call as “newly accelerated” naming it in terms of rhythm; hence,

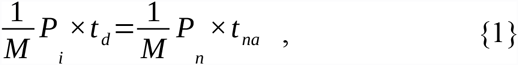

where _i_ – initial, _d_ – decelerated, _n_ – new, _na_ – newly accelerated.

The value of deformation *L*_2_ produces systolic pressure which overcomes elevated resistance ∑*R*:

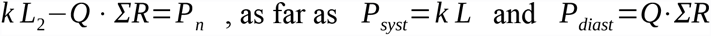

Hence, taking into account that 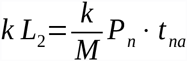 we use the right part for substitution:

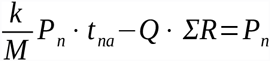, and after transforming we obtain 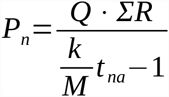.

Then we substitute this formula to {1} and reach the final expression:

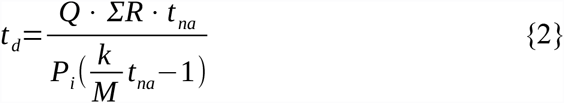

Consequently we must know beforehand what is the value of rhythm which is “newly accelerated” (*t*_*na*_), i.e., we must know the value of rhythm which is ought to be restored, and, as the way out, we must introduce (postulate) the concept of normal heart rate. (Certainly the normal rhythm may be chosen according to stipulation because there is no constant numerical value of *t*_*na*_ in {2}, i.e., our abstract calculations can be related to any rhythm considered as normal.) Therefore, we have postulated that among three parameters that are responsible for equilibrium (the deviator, point of balance and the counterbalance) the one is automatically reversible, i.e., it always recurs to its normal level, whereas two other parameters, -resistance, ∑*R*, (producing deviation) and venous pressure, *P*, - conserve their changed levels (or, better to say, conserve the equipoise achieved by the changed levels). It is clear that ∑*R* is the source of imbalance, i.e., the deviator; consequently, it is necessary to have at disposal: a point of balance and a counterbalance. Postulation of recurrence to normal rhythm is the introduction of the point of balance; respectively, venous pressure can play the role of counterbalance.

Formula {2} was deduced for the description of process of increasing of arterial resistance but evidently the process of decreasing is symmetrical with the process of increasing and, respectively, we must change indexes taking into consideration that now we need more general ones. Thus, index d (decelerated) must be replaced by ch (changed, i.e., decelerated or accelerated due to the rise or drop of the resistance) and index na (newly accelerated) must be replaced by N (normal, because we postulated the restoration of normal rhythm after elevation or decreasing of the venous pressure). Index _i_ (initial) at *P*_*i*_ remains the same because, in general case, the level of venous pressure that participates at formula {2} is the level registered before the forthcoming change of the resistance, i.e., *P*_N_ is the particular case of *P*_*i*_. So, we are to rewrite {2}:

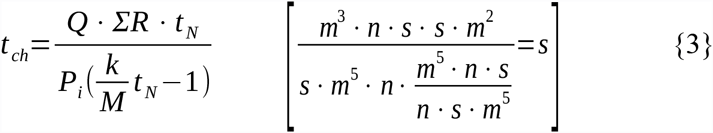

We have stopped tracing the reaction of the system in response to minimal elevation of the resistance (∆*R* = 10) when we have stumbled upon the question: what is the new value of rhythm (decelerated rhythm)? Now we can get the answer. Carotid reflex, firstly, detects the resistance by means of measuring of *P*_*diast*_ and in parallel information about ∑*R* comes directly from the muscles of arterial resistive vessels, - the pair which is necessary for calculation of volume flow rate (the problem of discrepancy between *P*_*diast*_ and ∑*R* will be scrutinized below). Secondly, carotid reflex, - or some “calculative center”, - solves the equation {3}, substituting *t*_N_ = 6, ∑*R* = 40 and *P*_*i*_ = *P*_N_ = 6 into {3}; hence, the value of elongated diastole appears and let it be denoted as *t*_*shift*_ because the change of duration of diastole takes place when the level of venous pressure is not changed yet, i.e., we observe the shift of the operating cell along the same line (in the table of **Fig.1**):

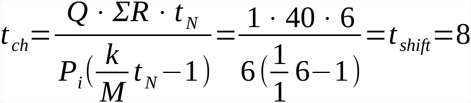

Thus, decelerated rhythm was found (*t*_*shift*_ = 8) and it must be multiplied by the value of the normal venous pressure *P*_i_ = *P*_*N*_ = 6 according to 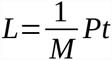 (and taking into consideration that 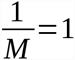); the product is the end-diastolic deformation *L* = 48 which generates systolic pressure numerically equal to *L*, i.e., *P*_*syst*_ = 48. As far as *P*_*diast*_ = Q • *ΣR* = 1 • 40 = 40 it is possible to find residual pressure (venous pressure) according to formula *P=P*_*syst*_*-Q· ΣR*. Hence, *P* = 48 – 40 = 8, i.e., the new *P*_*i*_ = 8. Horizontal thin non-doted arrow points to the left: the initial operating cell 6_p_×6_t_ passes on to the new operating cell 6_p_×8_t_. (Here and below: _p_ marks the line, i.e., *P*; and _t_ marks the column, i.e., *t*. When decimal fractures for denoting *t* and *P*_*i*_ appear we are in need to picture the calculated values of *t* and *P*_*i*_ inside the cells with whole indexes _t_ and _p_ which are closest, - or graphically convenient, - to calculated values of *t* and *P*_*i*_).

Bainbridge reflex, firstly, detects the level of venous pressure and, secondly, solves the same equation {3} that previously was solved by carotid reflex:

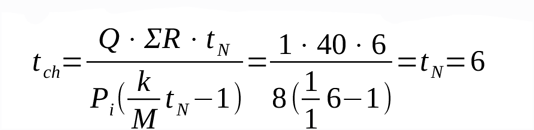

Therefore, Bainbridge reflex accelerates rhythm, i.e., it shortens the duration of diastole down to *t*_N_ = 6. In other words, Bainbridge reflex makes normal heart rate recurred: thin non-doted arrow points upwards and to the right, - i.e., the operating cell 6_p_×8_t_ passes on to the cell 8_p_×6_t_. Now circulation is stabilized in the respect of heart rate (the normal pulse is restored according to postulated condition that the system must be aware in advance about the value of rhythm the system must be returned to) but other measurable parameters are changed: the resistance (and *P*_*diast*_, respectively) is elevated from 30 up to 40, *P*_*syst*_ is increased from 36 up to 48 and the venous pressure is also increased from 6 up to 8. Thus, we now observe new equilibrium that has replaced the previous one which was poised within normal parameters only; this new equilibrium demonstrates the existence of two steps of searching the balance: 1) the violation of the balance by elevated resistance (the deviator) that makes the system to react immediately and restore equilibrium by changing of the point of balance (*t*_N_ = 6 was changed to *t*_*shift*_ = 8) and it is the transitory step; 2) then the counterbalance appears (raised venous pressure) and it is factually the repeated violation of balance (violation of the transitory balance), i.e., raised venous pressure compels the system to restore the previous point of balance (*t*_N_ = 6).

The way backwards begins from the command to decrease the resistance, i.e., ∆*R* = −10; obviously the resistance will fall from ∑*R* = 40 to *R*_N_ = 30 and the muscles of resistive arteries will inform the pacemaker (or “calculative center” of the pacemaker) about the drop of the resistance. In parallel the baroreceptors of carotid reflex will detect the decline of diastolic pressure which coincides with the decreasing of the resistance (due to the fact that *Q = 1*) and “calculative center” will start to solve the equation {3} in order to find *t*_*ch*_ (more accurately, *t*_*shift*_) which, for the present instance, denotes the acceleration of rhythm:

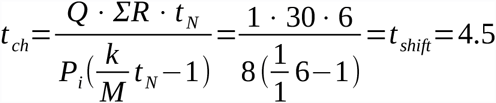

The operating cell is now 8_p_×4.5_t_ (located inside the cell 8_p_×5_t_ at **Fig.1**) – horizontal thin non-dotted arrow points to the right from the previous operating cell 8_p_×6_t_. The product, i.e., end-diastolic deformation, is *L* = 8 · 4.5 = 36. Diminished deformation generates the diminished systolic pressure which we consider numerically equal to *L*, i.e., *P*_*syst*_ = 36. This value of systolic pressure overcomes *P*_*diast*_ = *Q* · *R* _N_ = 1· 30 = 30; the residual pressure (venous pressure) is falling: *P=P*_*syst*_*-Q* · *ΣR* = *P*_*syst*_*-Q* · *R*_*N*_=36–1 · 30 = 6, and the decline of it will be detected by receptors of Bainbridge reflex (venous pressure descends from 8 to 6). Then Bainbridge reflex solves the equation {3} in order to find how intensively the rhythm must be decelerated:

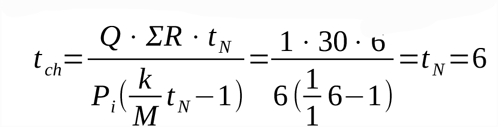

Thin non-dotted arrow points downwards and to the left; it indicates that the operating cell is again the cell 6_p_×6_t_; the value of *P*_*i*_ is now equal to 6, i.e., *P*_*i*_ = *P*_N_, and, therefore, circulation has restored its initial parameters (which were balanced before the deviation).

Two observations concerning this primary example of violation of balance and restoration of balance ought to be mentioned.

1. It seems that Bainbridge reflex does not need to calculate *t*_*ch*_ by means of equation {3} because Bainbridge reflex, as it seems, has the only one function - to normalize rhythm. What is it worth of measuring venous pressure then? The venous pressure can be elevated or can be declined as a consequence of perturbations at the arterial basin, i.e., it demonstrates the passive behavior. The objection is the following: both participants of regulation, - carotid reflex and Bainbridge reflex, - use the same formula {3} for calculating *t*_*ch*_ and, consequently, carotid reflex must be aware about venous pressure, *P*_*i*_, and, consequently, Bainbridge reflex must measure venous pressure and grant the data to carotid reflex; respectively, Bainbridge reflex must be aware of the resistance, ∑*R*. Besides, proper computing of *t*_*ch*_ claims the knowledge of the volume flow rate, *Q*, and we have already mentioned that *Q* can be found quite easily by means of comparison the information from baroreceptors (*P*_*diast*_) and information from the muscles of resistive arteries (direct information about ∑*R*). On the whole, we may emphasize that calculation of *t*_*ch*_ needs the existence of process of communication between arterial source of information and venous one.
2. When the arterial resistance is changed (and later such deviation will be superimposed on the forcing of volume flow rate) the role of Bainbridge reflex is to restore equipoise by means of normalizing of rhythm and it is really the restoration and nothing more. But the postulate of constancy of normal rhythm, as a point of balance, does not claim the conservation of some numerical value (once and forever) and permits to switch the mode of circulation choosing more rapid or more slow normal rhythm. Bainbridge reflex pretends to be the only one regulator of such switching, although such switching will inevitably generate new imbalance and, moreover, switching is not always feasible since the resistance may not permit. Evidently, the special role of Bainbridge reflex can hardly be fulfilled without calculations.

Thus, we have traced the reorganization of circulation caused by minimal rising of the resistance and further restoration of normal value of the resistance. **Fig.1** shows the examples of three more elevations of the resistance with increment ∆*R* = 10 and two declines of resistance with decrement ∆*R* = −10 (each step of rising or falling of resistance is accompanied by the recurrence to the initial state, i.e., by the recurrence to the previous step) and the reader can calculate by himself the parameters depicted in the operating cells of the table: *L –* end-diastolic deformation numerically equal to systolic pressure, *t –* duration of ventricular diastole (the value of *t*_N_ in column №*6* is omitted because it is always equal to 6, i.e., it is normal).

Now let us have a look at all the responses shown at **Fig.1** and determine the type of function between venous pressure, *P*_*i*_, and duration of ventricular diastole, *t*_*ch*_. We may represent formula {3} the following way: 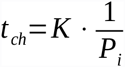, where 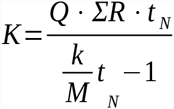 is the compound factor which corrects the value of *t*_*ch*_, i.e. *P*_*i*_ is a singular independent variable; therefore, the type of function is the function of inverse proportion (non-linear function). In that case, the variability of ∑*R, Q* and *t*_N_ will organize the linear correction of the value of *t*_*ch*_ because ∑*R, Q* and *t*_N_ belong to the numerator of *K* and the most complex is the correction accomplishing by ∑*R.* At **Fig.1** each line (*P_i_*) i.e., the argument, is accompanied by the value of the changed ∑*R* which is not shown; the increments of ∑*R* and *P*_*i*_ are steady: ∆*R = 10* and ∆*P*_*i*_ = 2 and it illustrates that ∑*R* is linked with *P*_*i*_ linearly. The diagram at **Fig.1** presents the duration of diastole as function and venous pressure as argument while the primary dependance of *t*_*ch*_ from the resistance, which we consider the correction, is omitted excepting the fact of positive or negative change of resistance (elevation or decreasing). Therefore, *P*_*i*_ (which is present at diagram as the independent variable) does not reflect the direction of the varying of ∑*R* and the opposite corrections by *+∑R* or *-∑R* (elevation or decreasing of the resistance) influence the function directly. We observe two inverse proportions: **Fig.1** shows two series of values of *t*_*ch*_ (more accurately, *t*_*shift*_) which frame two symmetrical graphs (visualized by operating cells in the table) and which look very much like hyperbolas. Symmetry axis is the column of operating cells (2, 4… 12)_p_×6_t_ where 6_t_ is the duration of normal ventricular diastole, i.e., symmetry axis denotes the equipoise at each level of *P*_*i*_ (the level caused by the change of ∑*R*). Other corrections (or deviations, as we call it), -slower or more rapid *t*_N_ and the forced *Q*, - just change the configuration of the paired hyperbols.

**Fig.2** and **Fig.3** show additional deviations that are provoked by substitution of some other values of normal duration of ventricular diastole (normal rhythm) instead of *t*_N_ = 6 into formula {3}; the change of *t*_N_ entails the change of normal venous pressure, *P*_N_, because normal resistance, *R*_N_, is still equal to 30. In other words, when we have shifted the position of the point of balance we observe the counterbalance (venous pressure) has changed its value while the resistance (the deviator) remains the same. The graph from **Fig.1**, where deviations of the resistance is shown at *t*_N_ = 6, is supplemented to **Fig.2** and **Fig.3** for comparison.

**Fig.2.**
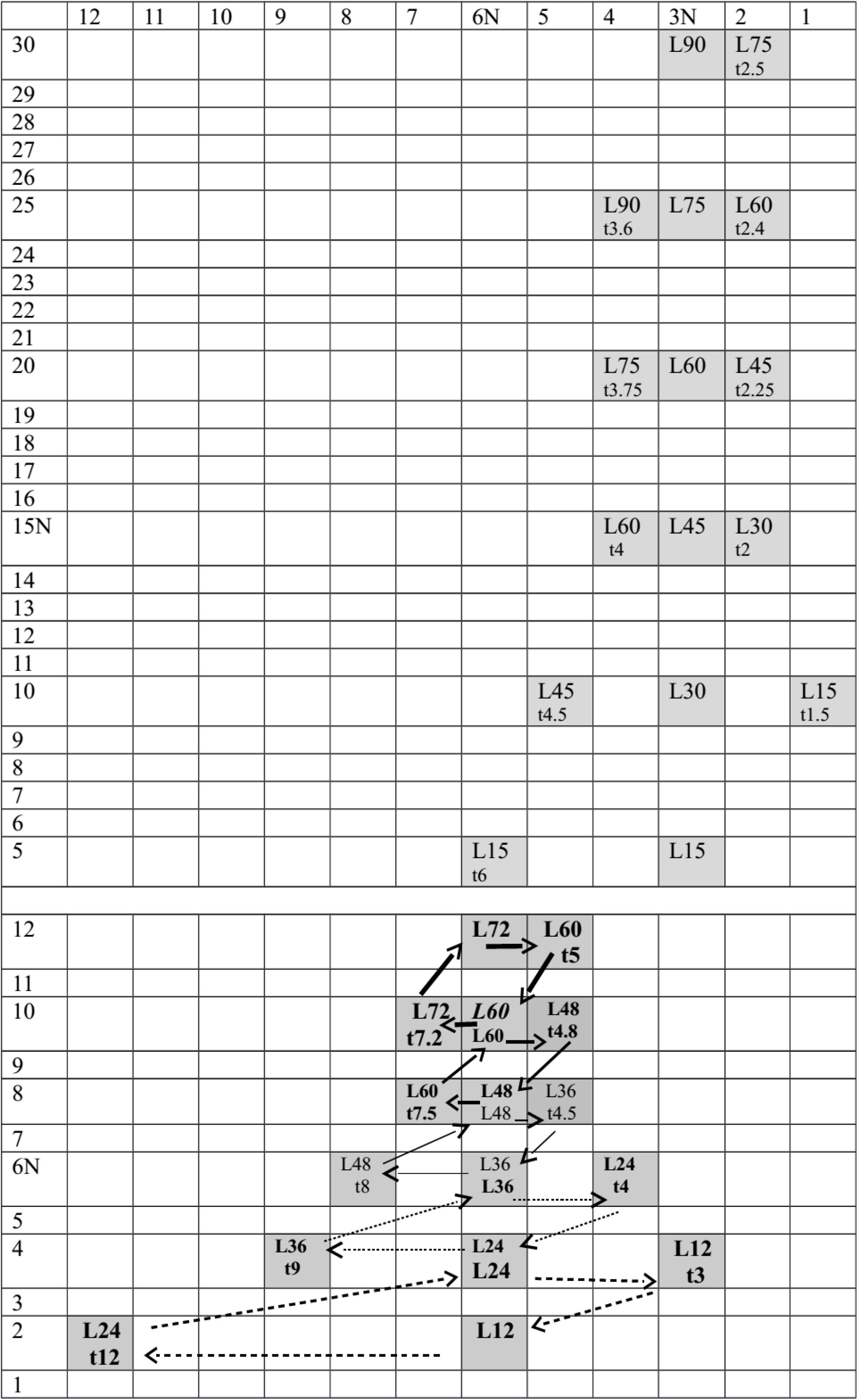
At the bottom there is a part repeated from **Fig.1**: the mode of circulation with normal values *t*_N_ = 6 and *P*_N_ = 6 (starting cell 6_p_×6_t_) which reacts to standard scale of variations of resistance. The upper part of the picture shows the mode of circulation with tachy-rhythm as normal (normal duration of ventricular diastole *t*_N_ = 3) and *P*_N_ = 15 (starting cell 15_p_×3_t_); the reaction to standard scale of variations of ∑*R* (arrows are omitted) is the following: diapason of venous pressures is broadened and shifted towards high values, diapason of rhythms (durations of ventricular diastole) is narrowed.

**Fig.3.**
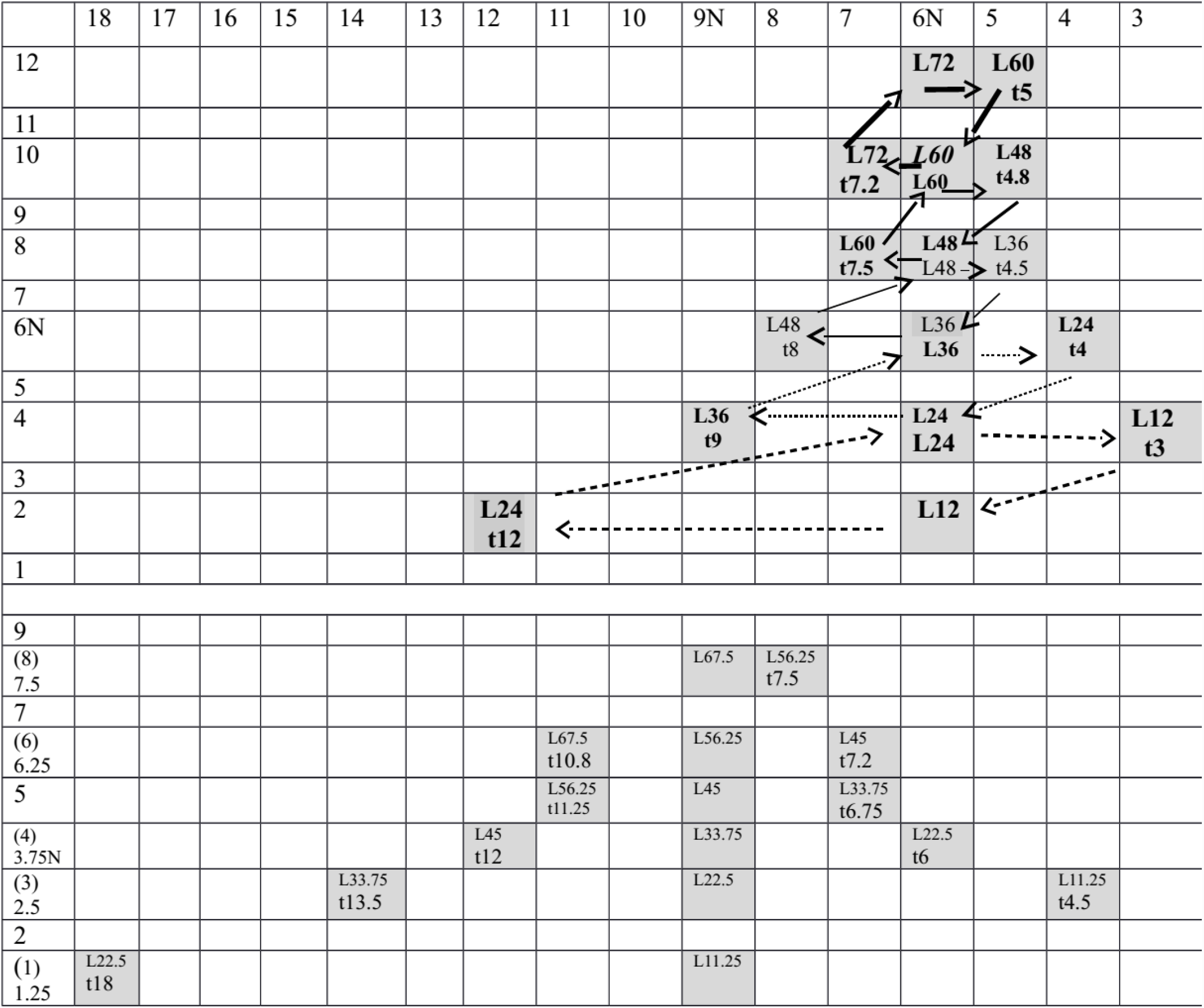
The picture is analogous to **Fig.2:** here the example for comparison from **Fig.1** is situated at the upper part of the picture; the mode of circulation with brady-rhythm as normal is placed at the bottom: *t*_N_ = 9, *P*_N_ = 3.75 (starting cell 3.75_p_×9_t_); the reaction to standard scale of variations of **∑***R* (arrows are omitted) is the following: diapason of venous pressures is narrowed and shifted towards low values, diapason of rhythms (durations of ventricular diastole) is broadened.

At **Fig.2** the new normal rhythm is more rapid, i.e., the duration of ventricular diastole is shorter and it is equal to 3 (3N at abscissa); the calculation of equipoise for standard normal resistance *R*_N_ = 30 results in changing of values of normal end-diastolic deformation of ventricle (*L* = 45) and normal venous pressure (*P*_N_ = 15 at ordinate). The regime of deviating of the system is the same as the initial one (which is shown near as a duplicate of graph from **Fig.1**), so we decided not to overload the construction with arrows indicating each step of deviation and further restoration of equilibrium, - we only marked out the operating cells of the table (this is true for **Fig.3** either). There are also two symmetrical hyperbolic graphs but general configuration is more stretched and compressed by the sides; it means that operating range of rhythms is much narrower than the one at the example shown for the comparison below, but operating range of venous pressure is broader. The parameter that mostly changed is venous pressure: *P*_*i*_ = 70 at the highest point comparing to *P*_*i*_ = 12 at the highest point at the picture below; therefore, circulation that is based on rapid heart rate maintains very high levels of venous pressure when the resistance is growing. Let us call this mode of regulation as tachy-mode and the initial one (**Fig.1** and the lower part of **Fig.2**) as norm-mode.

**Fig.3** presents the case with normal rhythm which is slower; the value of normal duration of diastole is *t*_N_ = 9, i.e., it is longer then *t*_N_ = 6 (norm-mode) and much longer then *t*_N_ = 3 (tachy-mode) and, respectively, this mode of circulation can be called - the brady-mode). New equilibrium, - while normal resistance is conserved (*R*_N_ = 30), - results in change of value of normal venous pressure and value of normal end-diastolic volume: *P*_N_ = 3.75 and *L* = 33.75, respectively. Configuration of the analogous symmetrical pair of hyperbolic graphs is flattened comparing to the configuration shown in the picture above (example with *t*_N_ = 6); it means that operating range of rhythms is broader but operating range of venous pressure is narrower and the latter is located at the low-pressure region. Therefore, this mode of circulation (where bradycardia was chosen as normal rhythm) is energetically favorable because such circulation can overcome standard growth of resistance exploiting low pressures.

Are there no advantages of tachy-mode at all? There is one significant advantage of tachy-mode that brady-mode lacks. It concerns the stability of the system when it reacts upon standard deviations (rising and falling of the resistance); the difference between tachy-mode and brady-mode can be easily illustrated by configurations of symmetrical hyperbolas: stretched (and compressed to the axis) hyperbolas of tachy-mode denote that decelerations and accelerations of rhythm are minimal, i.e., the system fluctuates within small amplitude of rhythms; on the contrary, flattened configurations of hyperbolas of brady-mode show that the system needs broad diapason of acceleration and deceleration of rhythm, i.e., the reorganization of the system (the response upon deviation) is in need of the enormous fluctuation of rhythm.

It is pertinent to suppose that the optimal variant is the following: to combine the both advantages (low pressures of brady-mode and high rhythmical stability of tachy-mode). As far as it is impossible to do simultaneously the question is the next: what part of brady-mode, - what levels of venous pressure (ordinate), - are accompanied by highest amplitude of fluctuation of duration of diastole (abscissa)? Obviously it is the level below the normal value of venous pressure (below 3.75N) when the branches of hyperbolas are most flattened.

Let us track out the case of brady-mode regulation (**Fig.3**, lower part) when the resistance decreased from 30N to 20: the duration of diastole shortened (*t*6) and the ventricle diminished its end-diastolic volume *L* = 3.75 • 6 = 22.5; the pulse difference now is equal to 2.5 (22.5 – 20) and this value denotes the new venous pressure; as far as venous pressure falls from normal value down to 2.5 the system will restore normal rhythm at that level of venous pressure (duration of diastole 9N), operating cell 2.5_p_×9_t_. The fluctuation of rhythm, in terms of duration of diastole, is equal to 3 (9 – 6). Imagine the system is programmed to defend itself from further possible high fluctuations of rhythm when the trend (first step) of falling of the resistance appears. The decision is to switch the brady-mode to the mode of regulation with more rapid normal rhythm – norm-mode (upper part of **Fig.3**) or tachy-mode. The transition from brady-mode to norm-mode is shorter, consequently, it is preferable. The goal is to convert the cell 2.5_p_×9_t_ into the nearest cell from norm-mode – with normal duration of diastole 6N (i.e., balanced like the cell 2.5_p_×9_t_) and with the same level of resistance (20); this level of resistance corresponds to the level of venous pressure equal to 4 of norm-mode (**Fig.3**, upper part), consequently, the goal is to turn to the cell 4_p_×6_t_. Hence, the equation is simple: *L*_?_ = *L*_24_, i.e., t_?_ • 2.5 = 6 • 4, and *t*_?_ = 9.6. Therefore, without any command from carotid reflex (the resistance does not change) the system decelerates rhythm and frames the cell 2.5_p_×9.6_t_ which originates the end-diastolic volume equal to 24 (2.5 • 9.6 = 24) instead of 22.5; the pulse difference rises from 2.5 to 4 (24 – 20), i.e., the venous pressure rises from 2.5 up to 4; then the system changes the mode of regulation (changes the value of normal duration of diastole of brady-mode, 9N, to the normal duration of diastole of norm-mode, 6N). The newly organized cell 4_p_×6_t_ belongs to norm-mode and, therefore, the future restoration of normal resistance (30N) will not claim to elongate diastole to 13.5 (the cell 2.5_p_×13.5_t_ at brady-mode) but only to 9 (the cell 4_p_×9_t_ at norm-mode) while searching of equipoise for *R* = 30.

The way back to brady-mode is just reverse and the initial command will be connected not with the falling of the resistance but with elevation of it. Diapasons of fluctuations of rhythm while balancing the system in response to elevation of the resistance are rather narrow at every mode of regulation. Certainly, the brady-mode is energetically preferable but nevertheless the modes with more rapid normal rhythm demonstrate smaller amplitudes of fluctuations and in cases when balancing of the system is more important there is no use to return to the brady-mode immediately. However, let us trace the switching of norm-mode with elevated resistance to the brady-mode with the same level of elevated resistance. Let us convert the cell 8_p_×6_t_ of norm-mode to the cell 5_p_×9_t_ of brady-mode; the latter is chosen because the level of resistance must be the same as it is at the cell 8_p_×6_t_ (*R* = 40) (**Fig.3**). The equation is the following: *L*_?_ = *L*_45_, i.e., *t*_?_ • 8 = 9 • 5, and *t*_?_ = 5.625. Therefore, without any command from carotid reflex (the resistance does not change) the system accelerates rhythm and frames the cell 8_p_×5.625_t_ which originates the end-diastolic volume equal to 45 instead of 48 (cell 8_p_×6_t_); the pulse difference falls from 8 (48 – 40) to 5 (45 – 40), i.e., the venous pressure diminishes from 8 to 5; then the system changes the mode of regulation (changes the value of normal duration of diastole of norm-mode, 6N, to the normal duration of diastole of brady-mode, 9N). The newly organized cell 5_p_×9_t_ belongs to brady-mode and, therefore, the system now continue overcoming the elevated resistance by means of systolic pressure (numerically equalized to systolic volume and end-diastolic volume) equal to 45 (*L*45) and venous pressure equal to 5 – instead of systolic volume equal to 48 (*L*48) and venous pressure equal to 8, respectively.

The important conclusion ensues. Initially the basic mode of the response to elevation of the resistance is brady-mode due to its energetic economy (i.e., circulation will chose the slowest possible rhythm as normal and this lower limit depends on some extra-circulatory reasons); but the elevation and further restoration of the resistance within brady-mode contains the trend to find oneself in disequilibrium and the system converts brady-mode into the mode with more rapid normal rhythm in order to prevent high amplitude of fluctuations of rhythm typical for brady-mode, i.e., the modes with rapid normal rhythm must be chosen when the system is in need of more high level of keeping the balance. The choice between energetic economy and stability of equilibrium forms the floating normal rhythm. Operating range of brady-rhythms is reduced to minimum and is preferable for steady state of circulation. Operating range of tachy-rhythms is much wider because the rhythmical response to significant deviations of resistance (rising of it and further restoration of it) lacks the danger of the very slow rhythms which can appear abruptly according to configuration of hyperbolas of brady-mode (but high levels of venous pressure which accompany the tachy-mode are not energetically economical).

Thus, let us have a look at formula {3} again. At first we have varied parameter ∑*R* and have obtained two symmetrical hyperbolic curves as the response to the deviation of the system; at that case, *t*_N_ and *Q* were considered constant. Coefficients *M* and *k* are considered invariable at this model. *M* is a coefficient of dynamic viscosity of relaxing myocardium; *M* lacks variability until the description of some mechanism which prevents rupture of extremely dilated ventricle improves the model; at present model such mechanism is omitted. Coefficient *k* is assumed equal to 1 (as simplification) and it defines the linearity that exists, firstly, between end-diastolic volume and systolic volume (Frank-Starling law) and, secondly, between systolic volume and systolic pressure (the latter bond, we repeat, is taken for granted now as numerical equalization of these two parameters but this linearity is not unfounded due to reference [4]; in that way, we may speak about invariability of *k*). The next step has included variation of the parameter *t*_N_ – but simultaneously the parameter ∑*R* has been repeating its previous varying, - and, as the result, we have found two different modes of circulation which contrast significantly while overcoming standard stepwise changes of the resistance. The only one parameter from formula {3} that has not been undergone to procedure of perturbation is the volume flow rate, *Q.* Previously we considered *Q* constant and equal to 1 keeping in mind that stability of cell metabolism provides the feedback for maintaining stability of *Q*; we are not going to specify this feedback but imaginary intensification of metabolism, - due to physical exertions, for instance, - must force the volume flow rate of blood. So it is quite reasonable to raise the value of *Q* and trace the consequences; for this aim *Q* will be intensified in the form of one-step forced value of *Q.* New value of *Q* will be applied to the two diametrically opposite modes of circulation which have been revealed while varying of the normal heart rate, i.e., to the tachy-mode and to the brady-mode. The new value of *Q* was chosen 1.2 instead of 1 for the convenience of visual proof only. We may remind that the information about intensification of volume flow rate is a calculable value: when the data coming from carotid baroreceptors does not coincide with data coming from muscles of resistive arteries the value of volume flow rate differs from 1. As far as *P*_*diast*_ = *Q · ΣR*, - i.e., *P*_*diast*_ is the information taken from carotid baroreceptors and *ΣR* is the information taken from muscles of resistive arteries, - the equation 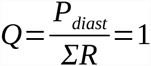 determines the case when two sources give identical information; if the fracture is equal to 1.2, for instance, it means that the discrepancy was caused by the command (from some upper level of regulation) to overstate *P*_*diast*_ deliberately and, obviously, such a command has a goal to force the volume flow rate.

The final part of the article includes the two examples concerning triple deviation; firstly, we have chosen most contrast modes of circulation (tachy-mode and brady-mode); secondly, we have forced the volume flow rate and, thirdly, we have undergone the both modes of circulation (with forced *Q*) to three-stepped elevation and two-stepped decline of the resistance; we excluded the returns from the steps to its starting positions, - as it was carried out when we have varied only the resistance (**Fig.1**) and the resistance together with the change of normal value of rhythm (**Fig.2** and **Fig.3**), - because it would overload the procedure.

**Fig.4** shows the pattern with rapid rhythm taken from **Fig.2**; the point for the intrusion is the equilibrium based on normal (for tachy-mode) duration of diastole and normal (for tachy-mode) venous pressure, i.e., the operating cell is15_p_×3_t_ (*t*_N_ = 3, *P*_*i*_ = *P*_N_ = 15 when ∑*R = R*_N_ = 0 and *Q* = 1). The new equilibrium will not affect *R*_N_ = 30 for the beginning but will only force volume flow rate (*Q* = 1.2 instead of *Q* = 1); so we are to find the values of *t*_N_ and *P*_N_ for new equilibrium. First of all let us calculate the new value of duration of diastole that will appear after the increasing of *Q*; it can be calculated as *t*_*shift*_ (venous pressure remains the same but duration of diastole will be changed) by means of substitution of all the parameters of operating cell 15_p_×3_t_ into formula {3} with the excepting of *Q* = 1 which now is *Q* = 1.2:

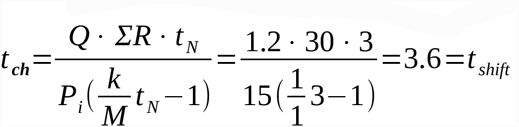

**Fig.4.**
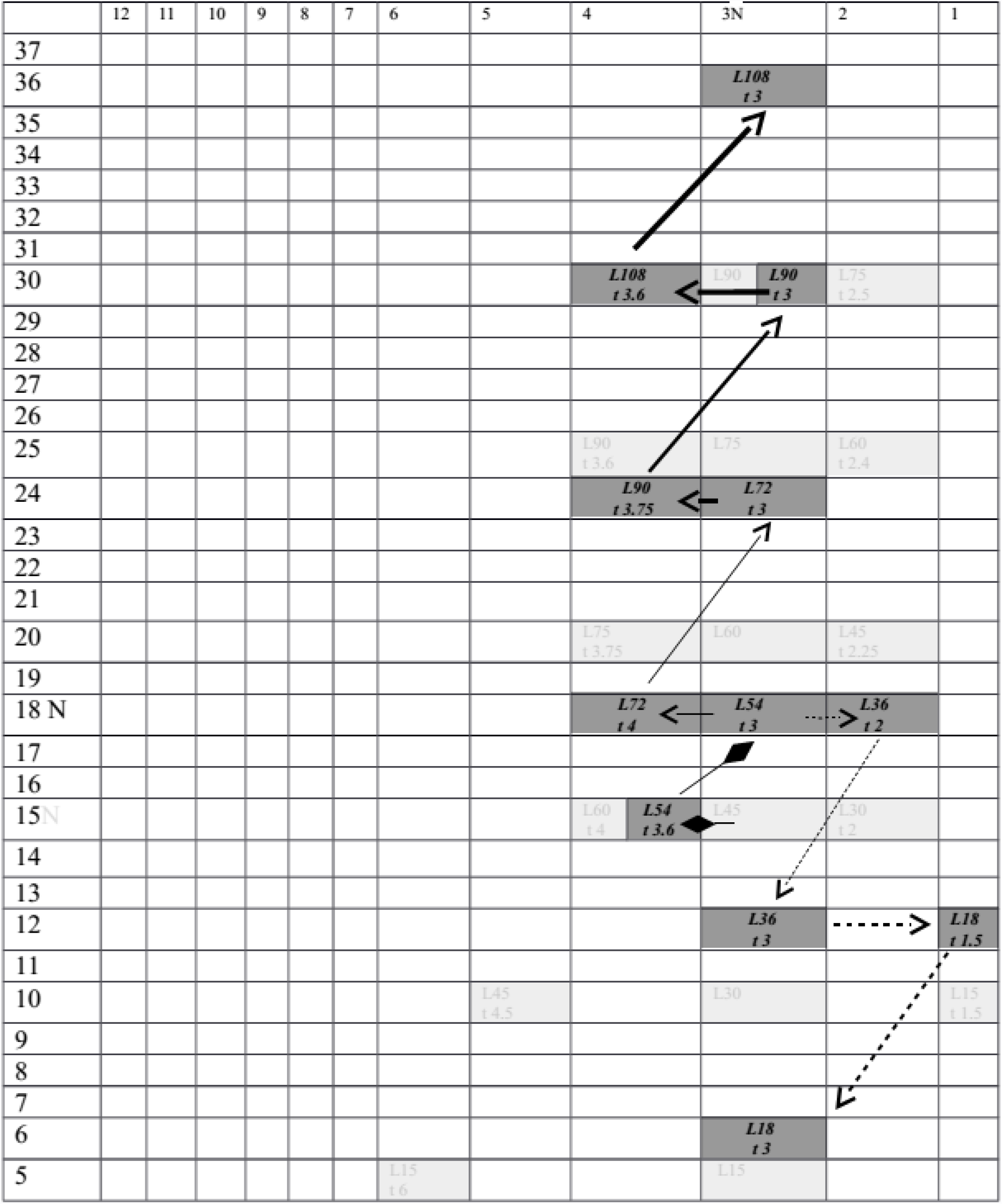
The background is the tachy-mode of circulation taken from **Fig.2**. The primary variation is the forcing of volume flow rate; the starting cell 15_p_×3_t_ which characterizes circulation at *R*_N_ and *Q* = 1 passes on to the cell 18_p_×3_t_ which balances circulation at *R*_N_ and *Q* = 1.2 (rhombic arrows). Next variation is the elevation and the decline of the resistance; first step of elevation of the resistance (∆*R* = 10, ∑*R* = 40) results in balancing of circulation at 24_p_×3_t_ (thin arrows). Three steps of elevation and two steps of decline of the resistance is represented.

Till now we have considered that information from carotid baroreceptors, – but not from muscles of resistive arteries, - was decisive, and may be it was partly true when *Q* = 1. Moreover, now when the resistance is not changed (∑*R* = *R*_N_ = 30, i.e., the information from muscles is not changed) the command to reorganize circulation can be originated from carotid baroreceptors only; evidently that it were not baroreceptors but “calculative center” which introduces the false information as if *P*_*diast*_ is already elevated (and, consequently, as if *Q* is already forced). The command results in calculation of *t*_*shift*_ and the prolonged diastole enlarges *L* and, hence, *P*_*syst*_ will be increased, - being numerically identical to *L*, - and, consequently, *P*_*diast*_ will be observed elevated because the resistance was not changed, - i.e., *Q* becomes forced the way it was planned by “calculative center”.

After multiplication *t*_*shift*_ = 3.6 by *P*_N_ = 15 we get the value of end-diastolic deformation *L* = 54; now operating cell is 15_p_×3.6_t_ (rhombic arrow directed to the left). Previously, when diastolic pressure numerically coincided with the resistance (*Q* = 1), we would subtract *P*_*diast*_ = *R*_N_ = 30 from *L* = 54 and would obtain the value of residual pressure (venous pressure) *P*_*i*_ = 24; but now *Q* = 1.2 and the correction is needed, consequently, *P*_*diast*_ = *Q* • *ΣR* = 1.2 • 30 = 36; hence, we subtract 36 from 54 and get *P*_*i*_ = 18. Bainbridge reflex detects the elevation of venous pressure (initially it was lower, *P*_N_ = 15) and begins to calculate by means of {3} the normal duration of the diastole, – i.e., Bainbridge reflex restores normal rhythm *t*_N_ = 3.

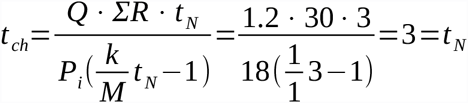

Now the operating cell is 18_p_×3_t_ (rhombic arrow directed upwards and to the right) and this cell presents the balanced circulation with forced volume flow rate *Q* = 1.2 and normal resistance *R*_N_ = 30; the indexes of the cell show that the normal (for tachy-mode) duration of diastole is the same but the level of venous pressure, - which is now considered normal for circulation with non-deviated resistance, - is *P*_N_ = 18, and that it is higher comparing to the level of venous pressure of the initial tachy-mode of circulation (*P*_N_ = 15). Therefore, this cell is the new equilibrium which denotes a new starting point for further elevation (or decline) of the resistance.

Now it is time to raise the resistance: ∆*R* = 10, ∑*R* = 40. The information from muscles of resistive arteries about the elevation of ∑*R*, - conducted along the pathway in parallel with the arc of carotid reflex, -results in deceleration of rhythm, i.e., this reflex is also programmed, like carotid reflex and Bainbridge reflex, that *t*_N_ = 3:

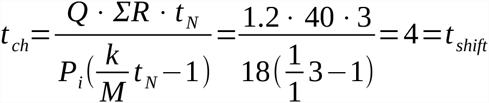

Hence, the operating cell is 18_p_×4_t_ (horizontal thin arrow directed to the left); end-diastolic deformation is now *L* = 18 • 4 = 72 and, therefore, operating cell is 18_p_×4_t_ (horizontal thin arrow directed to the left). *P*_*diast*_ = *Q* • *ΣR* = 1.2 • 40 = 48; then we subtract 48 from 72 and get *P*_*i*_ = 24. As far as venous pressure is detected (by Bainbridge reflex) elevated, - after the retardation of rhythm, - it means that equipoise is imbalanced and Bainbridge reflex is going to restore *t*_N_ = 3 by means of accelerating of rhythm and, consequently, calculates:

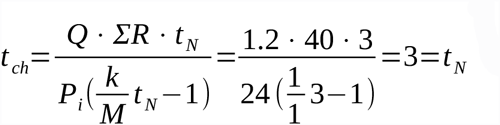

Now the operating cell is 24_p_×3_t_ (thin arrow directed upwards and to the right) and we may check up that it is a new equilibrium of circulation (with forced volume flow rate and elevated resistance): *L* = 24 • 3 = 72; *P*_*i*_ = 72 – 40 • 1.2 = 24.

Further reorganizations of circulation as the responses to step by step elevation (increment ∆*R* = 10) or decline (decrement ∆*R* = −10) of the resistance, - for tachy-mode of circulation, - can be easily calculated and traced with the help of **Fig.4**.

**Fig.5** visually describes the analogous forcing of volume flow rate and further rise and drop of the resistance – but variations are applied to the mode of circulation with slow rhythms (the pattern was taken from **Fig.3**). The starting point is the equilibrium with normal (for brady-mode) duration of ventricular diastole and normal (for brady-mode) venous pressure, - i.e., the operating cell is 3.75_p_×9_t_ (*t*_N_ = 9, *P*_*i*_ = *P*_N_ = 3.75 when ∑*R = R*_N_ = 30 and *Q* = 1). Carotid reflex (or, better to say, “the calculative center”) organizes the discrepancy between the data concerning *P*_*diast*_ and ∑*R* and inserts *Q* = 1.2 in formula {3}:

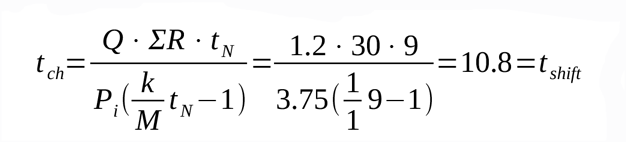

Now operating cell is 3.75_p_×10.8_t_ (rhombic arrow directed to the left). Multiplication of *t*_*shift*_ = 10.8 by *P*_N_ = 3.75 results in the value of end-diastolic deformation *L* = 40.5 which we numerically equate with systolic pressure; then it is necessary to find the value of *P*_*diast*_ which now differs from *R*_N_ = 30 due to the fact that *Q* = 1.2; consequently, *P*_*diast*_ = *Q* • *ΣR* = 1.2 • 30 = 36, and then we subtract *P*_*diast*_ = 36 from the value of systolic pressure numerically equal to *L* = 40.5 and get the value of residual pressure (venous pressure) *P*_*i*_ = 4.5. Bainbridge reflex detects the elevation of venous pressure and begins to calculate by means of {3} the normal duration of the diastole, *t*_N_ = 9 (restoration of normal rhythm):

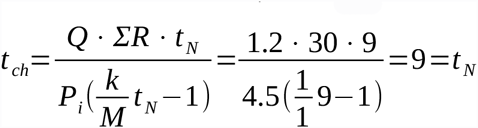

**Fig.5.**
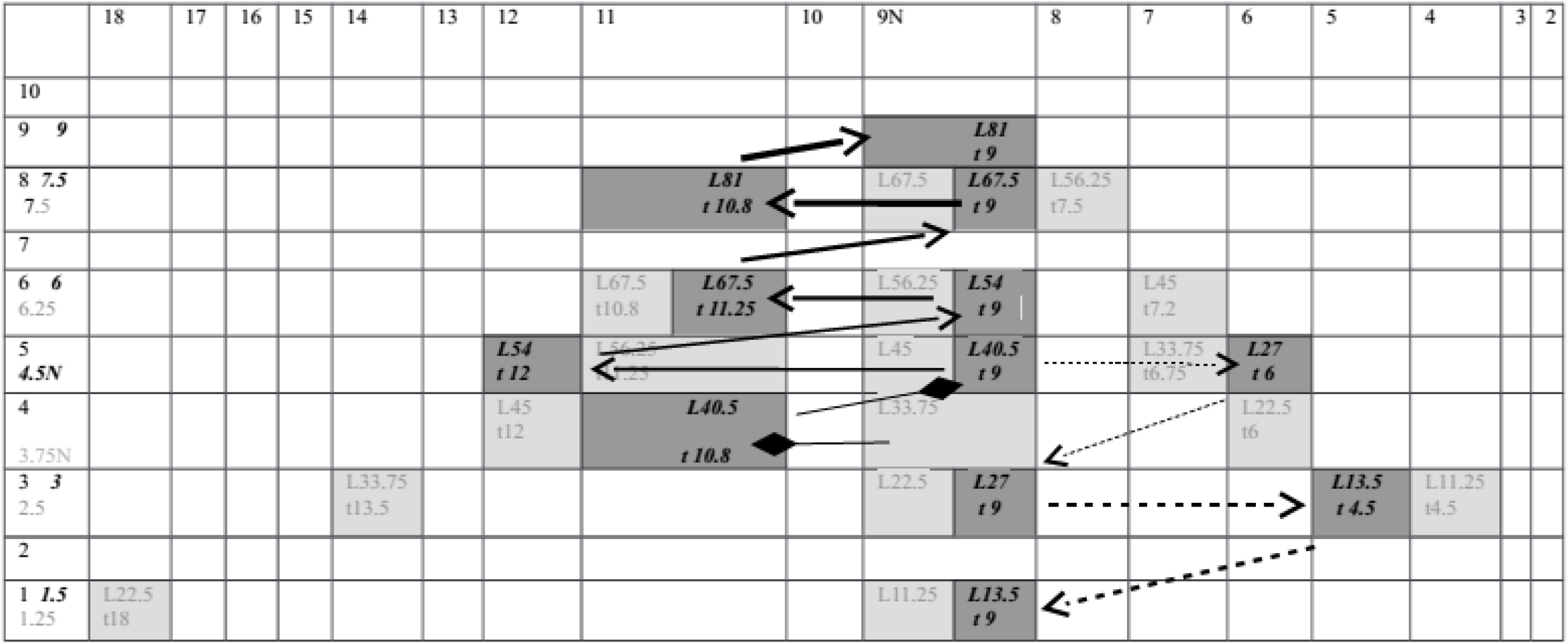
The background is the brady-mode of circulation taken from **Fig.3**. The primary variation is the forcing of volume flow rate; the starting cell 3.75_p_×9_t_ which characterizes circulation at *R*_N_ and *Q* = 1 passes on to the starting cell 4.5_p_×9_t_ which balances circulation at *R*_N_ and *Q* = 1.2 (rhombic arrows). Next variation is the one-step elevation of resistance (∆*R* = 10, ∑*R* = 40) which results in balancing of circulation at 6_p_×9_t_ (thin arrows). Three steps of elevation and two steps of decline of the resistance is represented.

The operating cell now is 4.5_p_×9_t_ (rhombic arrow directed upwards and to the right); this cell presents the balanced circulation with forced volume flow rate *Q* = 1.2 and normal resistance *R*_N_ = 30; the indexes of the cell show that the normal (for brady-mode) duration of diastole is the same but the level of venous pressure, - which is now considered normal, - is a new level (*P*_N_ = 18), and that it is higher comparing to the level of venous pressure of the initial brady-mode of circulation (*P*_N_ = 15). Therefore, this cell is the new equilibrium which denotes a new starting point for further elevation (or decline) of the resistance.

Now we are going to superimpose the minimal rise of resistance (∆*R* = 10, ∑*R* = 40) upon brady-mode of circulation with forced volume flow rate. The information from muscles of the resistive arteries about the elevation of ∑*R*, - conducted along the pathway in parallel with the arc of carotid reflex, - results in deceleration of rhythm, i.e., this reflex is also programmed, like carotid reflex and Bainbridge reflex, that *t*_N_ = 9:

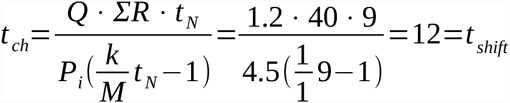

The operating cell now is 4.5_p_×12_t_ (horizontal thin arrow directed to the left). The end-diastolic deformation is *L* = 4.5 • 9 = 54 and it is numerically equal to systolic pressure; *P*_*diast*_ = *Q* • *ΣR* = 1.2 • 40 = 48; then we subtract 48 from 54 and get *P*_*i*_ = 6 and Bainbridge reflex succeeded in restoration of *t*_N_ = 9 by means of calculation:

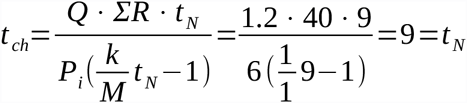

Now the operating cell is 6_p_×9_t_ (thin arrow directed upwards and to the right) and we may check up that it is a new equilibrium of circulation (with forced volume flow rate and elevated resistance): *L* = 6 • 9 = 54; *P*_*i*_ = 54 – 40 • 1.2 = 6.

Further reorganizations of circulation as the responses to step by step elevation (increment ∆*R* = 10) or decline (decrement ∆*R* = -10) of the resistance, - for brady-mode of circulation, - can be easily calculated and traced with the help of **Fig.5**.

What difference is evident when comparing tachy-mode (**Fig.4**) and brady-mode (**Fig.5**) of circulation, although both were undergone to standard forcing of *Q* and further elevation of the resistance? Tachy-mode starts with rapid rhythm (*t*_N_ = 3) and high level of venous pressure (*P*_N_ = 15); tachy-mode reacts upon the intensification of regime of circulation by means of significant rising of venous pressure (*P*_*max*_ – *P*_N_ = 36 – 15 = 21) and insignificant amplitude of fluctuation of rhythm (deceleration: *t*_*max*_ - t _N_ = 4 – 3 = 1). Brady-mode has another starting point: slow rhythm (*t*_N_ =9) and low venous pressures (*P*_N_ = 3.75); brady-mode reacts with moderate amplitude of fluctuation of rhythm (deceleration: *t*_*max*_ - *t*_N_ = 12 – 9 = 3) and moderate rising of venous pressure (*P*_*max*_ - *P*_N_ = 3.75 – 9 = 5.25). Tachy-mode demonstrates 4-fold higher elevated venous pressure comparing to the elevation of venous pressure of brady-mode (21 : 5.25 = 4); brady-mode demonstrates 3-fold wider amplitude of fluctuation of rhythm comparing to the amplitude of fluctuation of rhythm demonstrated by tachy-mode (3 : 1 = 3). It means that tachy-mode has difficulties with venous pressure and brady-mode has difficulties with rhythm. What is it easier to overcome? Let us analyse what are these difficulties in the flesh. Firstly, we may add to analysis the values of overall diapasons of systolic pressure of tachy-mode (*P*_*syst max*_ – *P*_*syst N*_ = 108 – 45 = 63) and brady-mode (*P*_*syst max*_ – *P*_*syst N*_ = 81 – 33.75 = 47.25) as far as the value of end-diastolic volume *L* was assumed to be numerically equal to systolic pressure. Hydrodynamics with higher pressures spends more energy and, consequently, tachy-mode is not energetically favorable. As regards to brady-mode the demonstrative negative feature is 3-fold wider amplitude of fluctuation of rhythm comparing to tachy-mode; serious trend of disequilibrium can be expected if the circulation with the forced volume flow rate encounters with the falling of the resistance and further restoration of it (lower parts of hyperbolas). That is why the conclusion concerning the cases with forced volume flow rate can be referred to the one that has been made above: any deviation of the resistance does not trend towards disequilibrium if the system temporary transmits the mode of response from brady-mode (which is good for steady state or for the small deviations of resistance) to tachy-mode.

We may apply this knowledge (about the advantages and drawbacks of the different operating ranges of rhythms and pressures specific to the opposed modes of circulation) to the phenomenon which lacks explanation within any modern theory of circulation (so far as we know). It is just the observing phenomenon that the normal pulse, as one understands it while measuring it at conditions of the rest, is located asymmetrically between the range of rhythms which circulatory system is using for deceleration and the range of rhythms which circulatory system is using for acceleration; such normality (steady rhythm in conditions of rest) is much closer to bradycardia then to tachycardia and the operating range of brady-rhythms is much narrower than the operating range of tachy-rhythms. If we admit the coincidence of the conclusion with the real phenomenon we must admit the whole chain of testimony including the main statement the testimony has been proceeded from. The main statement is that the relaxing ventricle imitates viscous deformation (deformation that has two independent variables, stress and time) and this statement has permitted to deduce the equation which successfully balances one-loop circulation.

